# Modeling the Role of the Striatum in Non-Stationary Bandit Tasks

**DOI:** 10.1101/196543

**Authors:** Sabyasachi Shivkumar, V. Srinivasa Chakravarthy, Nicolas P. Rougier

## Abstract

Decision making in non-stationary and stochastic environments can be interpreted as a variant of non-stationary multi armed bandit task where the optimal decision requires identification of the current context. We formalize the problem using a Bayesian approach taking biological constraints into account (limited memory) that allow us to define a sub-optimal theoretical model. From this theoretical model, we derive a biological model of the striatum based on its micro-anatomy that is able to learn state and action representations. We show that this model matches the theoretical model for low stochasticity in the environment and could be considered as a neural implementation of the theoretical model. Both models are tested on non-stationary multi-armed bandit task and compared to animal performances.

**Author Summary:** Decision making in changing environments requires knowledge of the current context in order to adapt the response to the environment. Such context identification is based on the recent history of actions and their outcome: when some action used to be rewarded but is not anymore, it might be a sign of a context change. An ideal observer with infinite memory could optimally estimate the current context and act accordingly. Taking biological constraint into account, we show that a model of the striatum, which is the largest nucleus of the basal ganglia, can solve the task in a sub-optimal way as it has been shown to be the case in rats in a T-maze task.

## 1 Introduction

In their day to day life, animals face two kinds of uncertainty. One that is expected and one that is not. For example, if an animal try to jump to catch a prey, there is a known risk of failure such that the prey can eventually evade the predator. Depending on the direction of the wind and the terrain topology, the prey can smell the predator and this latter has thus to decide what is the best distance to jump. Those parameters represent the context from which the predator has to adapt its behavior. Failing to do so means death in the long term. The difficulty is to appreciate the direction of the wind that may locally vary because of terrain topology and local turbulences. A variation can be transient, meaning the direction of the wind has not really changed, but such variation can be also the sign of a more global change in the direction of the wind. If the predator fails to characterize such context, a jumping distance that used to be safe in one given context won’t be in another one. Ultimately, this could mean a failure at catching the prey.

Earlier studies have categorized and named such randomness into expected and unexpected uncertainty (Yu & Dayan, 2003). Expected uncertainty refers to the variability in the different parameters of the environmental model constructed by the agent (since we use agents to model animals performing reward based tasks, we use the terms animal and agent interchangeably). A typical example of this is a stochastic reward distribution when the agent is performing a reward based learning task. Standard reinforcement learning models have been used to tackle problems with expected uncertainty (Kaelbling, Liftman, & Moore, 1996; Sutton & Barto, 1998). Unexpected uncertainty on the other hand refers to the case where there is a consistent difference between observations and the prediction by the agent. This could occur for example when there is a change in the environment (non-stationary environment). Specialised reinforcement learning models like modular reinforcement learning can identify the context of the environment and are therefore successful in solving such tasks. In this work, we are mostly interested in the role of the striatum in reward based tasks that involve both expected and unexpected uncertainty.

A lot of results tends to identify the basal ganglia (BG) as a key player in reward based learning tasks and model them as a reinforcement learning (RL) engine (Joel, Niv, & Ruppin, 2002; Chakravarthy, Joseph, & Bapi, 2010). Furthermore, striatum, which is a major component of the BG, has a rich microcircuitry consisting in central structures called striosomes, and matrisomes surrounding the striosomes (Graybiel, Flaherty, & Giménez-Amaya, 1991). The striatum is believed to form representations of state and action space used for performing RL tasks (Charpier & Deniau, 1997). The striosomes are believed to map the state space (Wilson, Takahashi, Schoenbaum, & Niv, 2014) while the matrisomes are believed to map the action space (Flaherty & Graybiel, 1994) based on their differential cortical projections. In addition, the striatum has reciprocal projections to both the Ventral Tegmental Area (VTA) and the Substantia Nigra pars compacta (SNc). It receives reward prediction error from these midbrain nuclei and uses it to map the developed representations to state (Granger, 2006) and action values (Seo, Lee, & Averbeck, 2012) which are used for action selection. The striatum has also been hypothesized to perform context dependent tasks by mapping different contexts to different striatal modules (Amemori, Gibb, & Graybiel, 2011).

In this article, we focus on stochastic and non-stationary tasks and we develop both theoretical and biologically plausible models to solve them. After formalizing the task definition, we derive an ideal Bayesian model for solving such tasks (Lloyd & Leslie, 2013). Considering some realistic task constraints on the model, we modified the theoretical Bayesian model to solve the problem iteratively. Following this idea, we develop a model of the striatum to handle these tasks. More precisely, we use a layered Self Organizing Map (Kohonen, 2001) architecture to model the striosomes and matrisomes as Strio-SOM and Matri-SOM where a single Strio-SOM neuron projects to the surrounding Matri-SOM neurons. The Strio-SOM and the Matri-SOM activity are mapped to compute state and action values, respectively and used for action selection. This striatal model is extended to a multi module based architecture to deal with multiple context paradigms. The biological plausibility imposes on the model limitations such as finite memory which is also incorporated into the theoretical model. Thus the theoretical model sets a bound on the expected performance for a probabilistic context dependent task. We show that the neural model is able to meet the optimal bound for low values of stochasticity in reward.

## 2 Materials and Methods

### 2.1 Non-Stationary Bandit Task

In the classical multi-armed bandit task (Auer, Cesa-Bianchi, & Fischer, 2002), at each time *t* ∈ {1,…, T}, an agent chooses an arm *a_t_* ∈ {1,…, *N*} according to a policy π and receives a reward *R_t_*(*a_t_*). The rewards 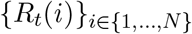 are drawn from a set of unknown distribution *p_i_* with expectation *µ*(*i*). The agent’s objective is to find a policy π such as to maximize the cumulated expected reward 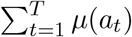. The optimal policy (oracle) π*\** consists in choosing *a_t_* = *i\** such that 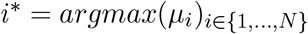.

In the non-stationary case, the rewards 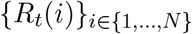 are drawn from a set of unknown and non-stationary distribution *p_i_*(*t*) that may vary over time. These variations can be continuous (Slivkins & Upfal, 2008) or abrupt (Garivier & Moulines, 2011; Besbes, Gur, & Zeevi, 2014; Raj & Kalyani, 2017). In this work, we‘ll consider only abruptly changing environments: the distribution of rewards remains stationary for a given period and a change occurs at an unknown time instants (breakpoints). The period between two breakpoints is called a context. In the general case, the number of breakpoints and the number of different context is unknown. In the present study however, we’ll restrict to the case *N* = 2 using a set of two different contexts (with several possible breakpoints alternating between the two contexts).

Such non-stationary bandit task can be related to some extent to a change-point detection problem that has been extensively studied in the context of controlled dynamic systems (Hinkley, 1970; Basseville & Nikiforov, 1993). However in the case of non-stationary bandit task, the agent is not a mere observer but an actor that may choose to pull this or that arm in order to sample the current context and gain information. The expected regret (difference between optimal reward and obtained reward) can be bound in both the stationary and non stationary bandit tasks. There exist some near optimal algorithms which aim to achieve this bound in stationary (Agrawal, 1995) and non-stationary bandit tasks (Garivier & Moulines, 2011; Besbes et al., 2014; Raj & Kalyani, 2017). While the policy proposed by these algorithms is optimal even in our problem for maximizing expected reward, they assume knowledge of certain parameters like T (length of context) which is not available to the agent. In addition, they cannot cope with biological constraints such as finite size memory and limited computational resources. This motivates the need for a biologically plausible optimal policy.

### 2.2 Bayesian formulation

We consider a non-stationary two-armed bandit task (arms *a*_1_ and *a*_2_) using two different contexts (*c*_1_ and *c*_2_) that can change at unknown times. In context *c*_1_, the optimal action is *a*_1_ and in context *c*_2_, the optimal action is *a*_2_. The case where the optimal action is the same in the two contexts is not studied here. The reward for the optimal arm is *R^+^* and the reward for the sub-optimal arm is *R*^−^, independently of the context. The reward distribution matrix is denfined as:

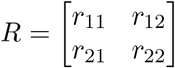

where *r_ij_* is the probability of getting *R^+^* when choosing arm *a_i_* in context *c_j_*. Reciprocally 1 — *r_ij_* is the probability of getting *R*^−^ when choosing arm *a_i_* in context *c_j_*.

Using Bayes theorem, and noticing that *P*(*c* = *c*_1_) *=* 1 — *P*(*c* = *c*_2_), we have:

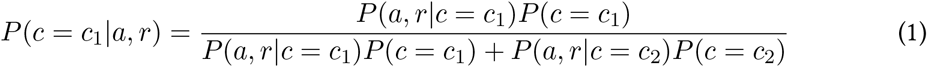

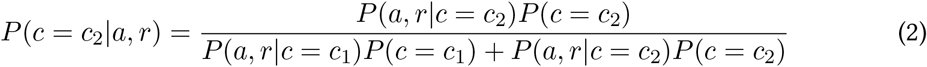

Considering a single trial and assuming we have no initial knowledge of the current context, we have *P*(*c* = *c*_1_) = *P*(*c* = *c*_2_) = 0.5 which allow us to rewrite equation (1) as:

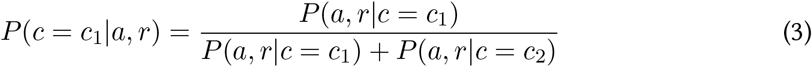

We can now extend this to multiple trials by keeping track of the history of action selection and rewards. We assume that our prior is stationary and remains constant across the trials. In other words, 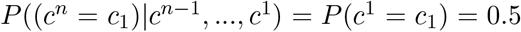, where *c^i^* is context in the *i^th^* trial. This is the case when the context can change every trial independent of the previous trials. Thus, at the *i^th^* trial, considering the chosen action *a^i^* and the reward r^i^, we have:

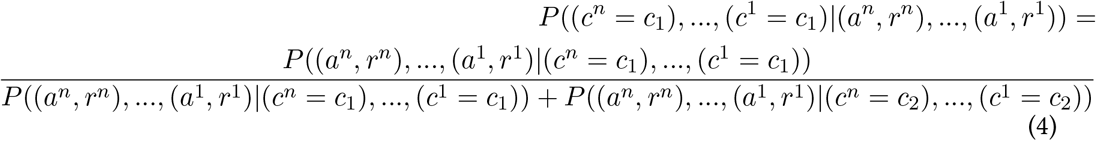

Since the action-reward pairs across trials are independent, equation (4) can be further simplified into:

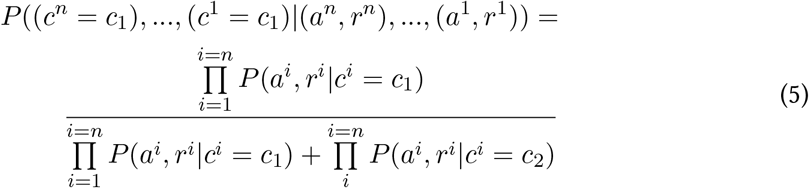

Equation (5) requires to keep track the full event history (action and reward). It can be simplified using a local average of the observed events (sliding window) using only the *τ* last plays as it has been proposed in the SW-UCB algorithm and proved to be almost rate-optimal in a minimax sense (Garivier & Moulines, 2011).

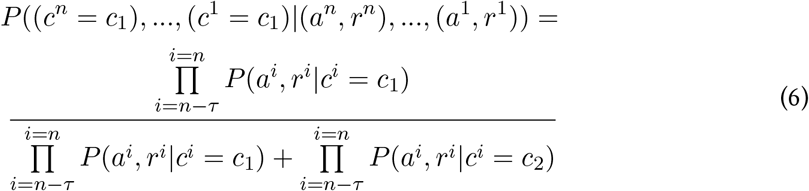

### 2.3 Theoretical Model

Even though we derived equation (6) to express the probability for a given context to be the actual context, we still need to estimate the different terms since they’re not accessible to the agent. Our next best option is thus to use an estimate of the current context denoted c from which we can derive a new estimate:

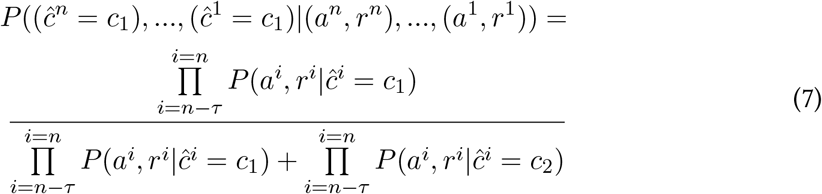

Using equation (7), we can now estimate the current context based on action and reward which writes:

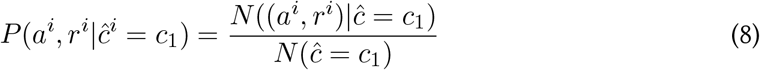

where 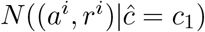 is the number of times the agent chose *a^i^* when it estimates the context as being *c*_1_ and get the reward *r^i^* and 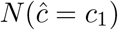 is the number of times the agent estimates the context as being *c*_1_. This expression was derived such that agent can estimate the context it is in by looking at the term 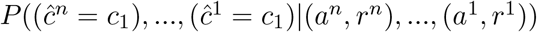.

However, in order to calculate this, we need terms that imply that the agent has to estimate the context and choose actions accordingly. There is thus an inherent circularity in the problem. To break this circularity, we have to solve the problem iteratively. We try to estimate the reward distribution function at trial number *t* and denote this as 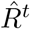. Furthermore, we keep track of another matrix 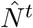 which counts the number of times the agent chose a particular action in a particular estimated context. These two matrices reads as follows:

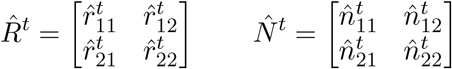

where 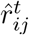 represents the estimated probability of getting a reward *R^+^* when choosing action *a_j_* in estimated context 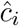 at trial *t*, 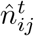 represents the number of times the agent chose action *a_i_* in estimated context 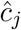 at trial *t*. For ease of notation, we define the likelihood to be 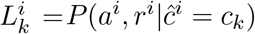 with *k* ∈ {1, 2}. Equation (7) now reads:

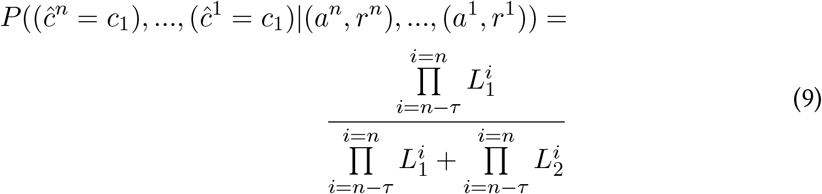

Assuming reward distributions are initially equal, we have:

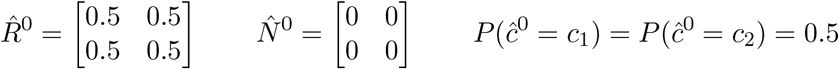

At trial *t*, the agent estimates the current context 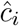 according to the estimate in previous trials and chooses the action (*a_j_*) according to equations:

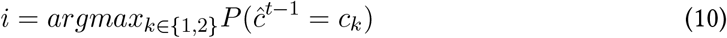

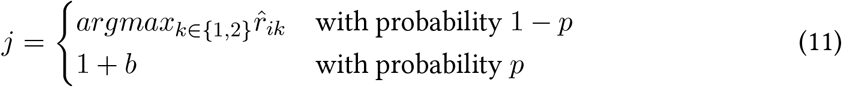

where *p* is the probability of exploration and *b* ~ *Bernoulli*(0.5). The exploration ensures that all the actions are sampled in the initial trials.

Based on the choice of 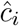 and *a_j_*, the agent can update the values of 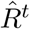 and 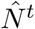 according to:

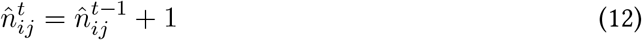

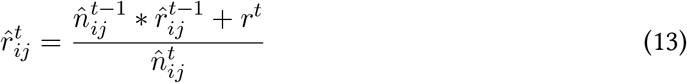

where 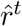 is the reward obtained at trial t.

Furthermore, since 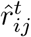 represents the estimated probability of getting a reward *R^+^* when choosing action *a_j_* in estimated context 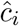 at trial *t*, 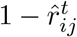 represents the estimated probability of getting a reward *R*^−^. Consequently 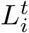 is given by:

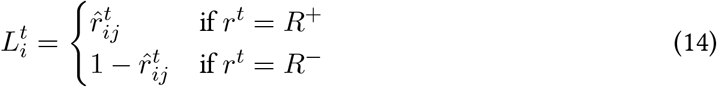

Finally, substituting values of equation (14) in equation (9) yields:

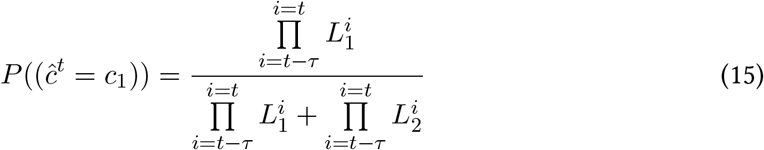

Equations (10) to (15) can now be used to formulate an algorithm in order to solve the nonstationary bandit task as shown in Figure 1.

**Figure 1.**
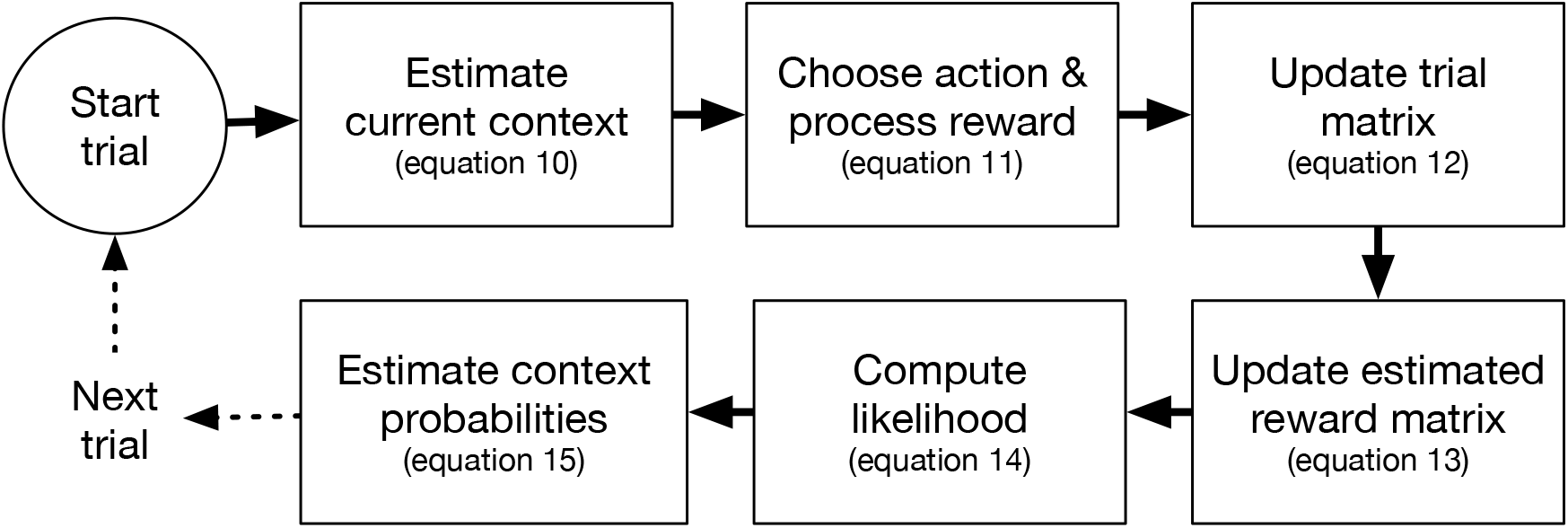
A synthesized view of the model showing the different steps necessary to solve a non stationary bandit task.

### 2.4 Neural Model (single module)

We proposed a theoretical model in the last section to solve non stationary bandit tasks. In this section we develop a biologically plausible model of the striatum for these tasks. This model is derived from an existing model of the basal ganglia proposed to solve non stationary problems (Shivkumar, Muralidharan, & Chakravarthy, 2017). The center-surround structures seen in the striatum are modeled using a layered SOM model. In a layered SOM model, each neuron in the center SOM layer projects to a secondary SOM layer.

The center layer in the striatal model is the Strio-SOM, which maps the state space and is believed to model the striosomes. The neurons in the Strio-SOM project to the Matri-SOM which maps the action space and is believed to model the matrisomes (Figure 2). Given *m*_1_×*n*_1_ neurons in the Strio-SOM and *m*_2_ × *n*_2_ neurons in the Matri-SOM, the weights of the Strio-SOM(*W*^S^) have dimension *m*_1_ × *n*_1_ × *dim*(*s*) where s is the state vector. Similarly, for an action vector a the weights of all the Matri-SOMs (*W^M^*) are of dimension *m*_1_ × *n*_1_ × *m*_2_ × *n*_2_ × *dim*(*a*) as each neuron in the Strio-SOM projects to a Matri-SOM. For a state input s, the activity for a neuron n in the Strio-SOM is given in (16).

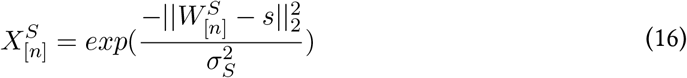

where [*n*] represents the spatial location of the neuron n and *σ_S_* controls the spread of the neuron activity. The complete activity of the Strio-SOM (*X^S^*) is the combination of individual activity of all the neurons. The neuron with the highest activity (”winner”) for a state s is denoted by 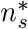

Similarly, for an action input a corresponding to a state s, the activity for a neuron n in the Matri-SOM is given in (17).

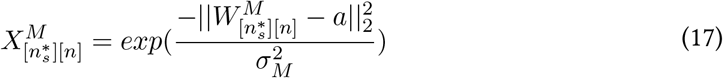

where *σ_M_* controls the spread of the neuron activity. The complete activity of the Matri-SOM corresponding to neuron 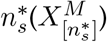 is the combination of individual activities of all the neurons in the Matri-SOM corresponding to 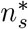. The neuron with the highest activity (”winner”) for an action a in state s is denoted as 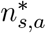. The weight of a neuron n in the Strio-SOM for a state input s is updated according to (18)

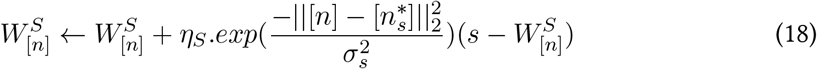

The weight of neuron n in the Matri-SOM for an action input a in a state s is updated according to (19).

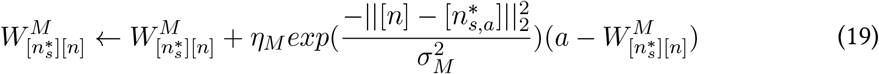

These representations can be used to evaluate the states and actions and guide the decision making process. The schematic of our striatal model to solve stochastic RL tasks is given in Figure 2.

**Figure 2.**
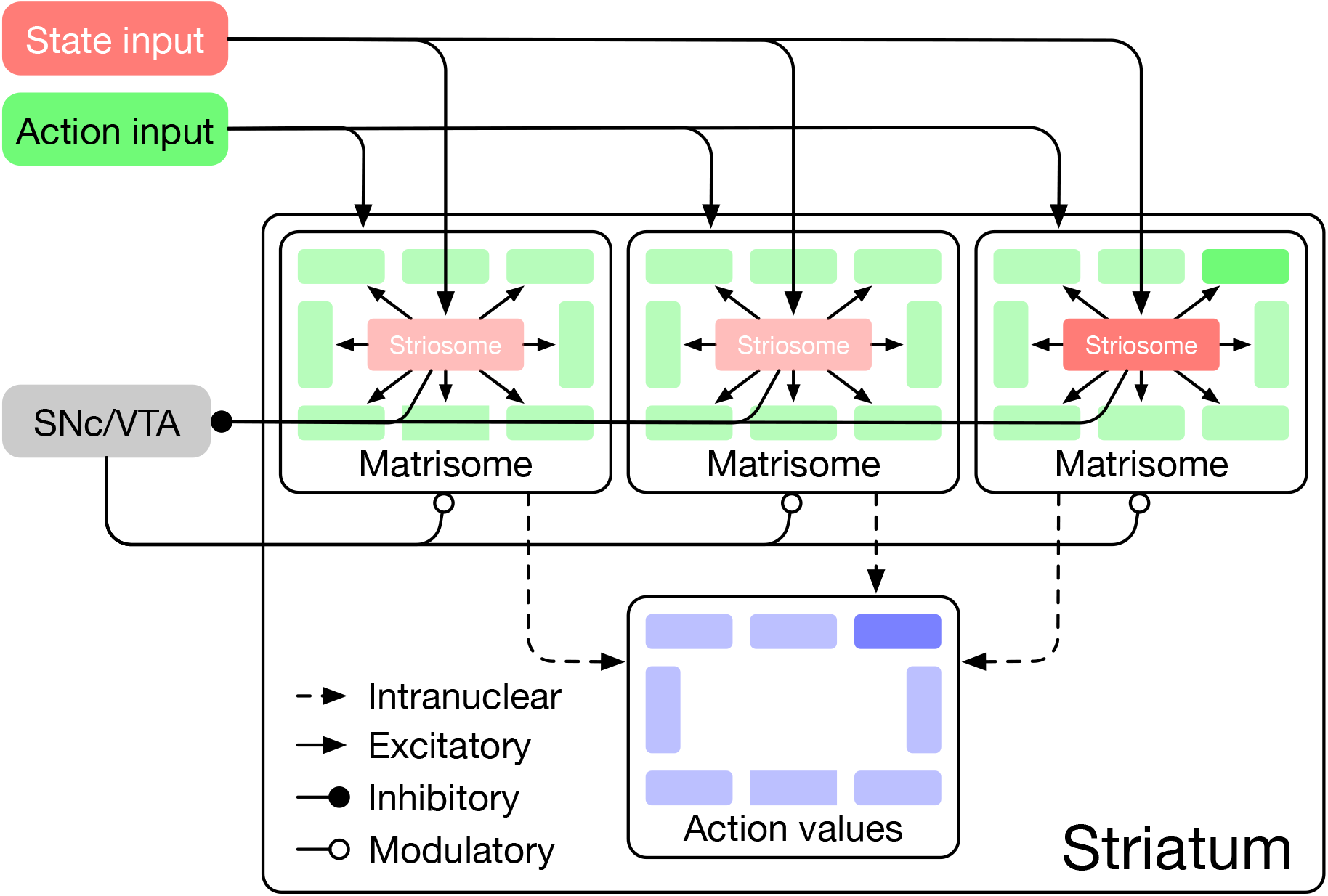
Schematic diagram of the Striatum model where the arrows indicate the connections and their types. The state input is mapped by the striosomes and the surrounding matrisomes map the actions possible in that state.

Let the agent performing the task be in state s. The striosome activity gives us the representation of the state in the striatum. This activity is modeled by the Strio-SOM as given in (16). Thus the activity is of dimension *m*_1_ × *n*_1_.

This activity of the Strio-SOM projects to the SNc and represents the value for the state s in our model (20). The Striatal-SNc 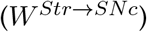 are trained using the signal from SNc which is representative of Temporal Difference (TD) error (*δ*) (21). The TD error is calculated as *δ* = *r +* γ*V*(*s′*) *— V*(*s*) where *s′* is the new state after taking action **a**, **r** is the reward obtained and *γ* is the discount factor.

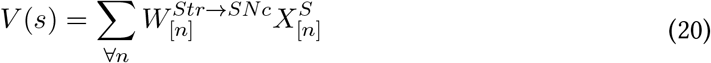

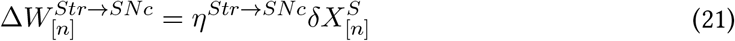

where V(s) represents the value for state s, 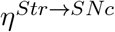 is the learning rate for 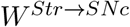.

The actions that can be performed in a state s are represented by the matrisome activity surrounding the striosome neuron for that state. This is given by the activity of the Matri-SOM corresponding to the neuron with the highest activity in the Strio-SOM 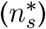 in our model. The activity of a Matri-SOM neuron for an action a is given in (17) and is of dimension *m*_2_ × *n*_2_.

The Matri-SOM activity X for action **a** is projected to the action value neurons as given in (22). If *n_a_* is the action value neuron for the action **a**, 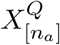, corresponds to the action value for **a** the action in the state **s** in our model. These connections are also trained using TD error as above and the update equation is given in (23)

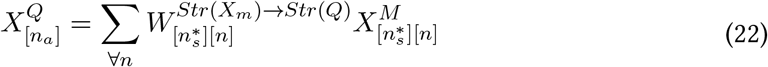

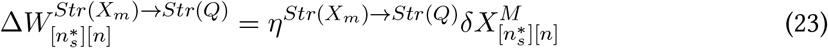

where *X^Q^* represents the activity of the action value neurons, 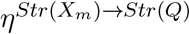 is the learning rate for 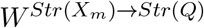.

The activity of the action value neurons are used for action selection by using a softmax policy in our model (24). We believe that this is carried out by the dynamics of the STN- GPe oscillations with the striatal action value neurons projecting to the GPe. This is further elaborated in the ‘Discussion’ section.

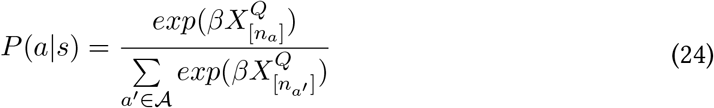

where *β* is the inverse temperature and 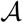 denotes the action set for the agent.

### 2.5 Neural Model (multi-modules)

The modular nature of the striatal anatomy has been proposed to be responsible for solving non stationary tasks using a modular RL framework (Shivkumar et al., 2017). In this method, the agent allocates separate modules to separate contexts. Each of the modules has its own copy of the environment in a particular context, represented by an environment feature signal (*ρ*). This copy is used to generate a responsibility signal, denoted by λ, which indicates how close the current context is to the one represented by the module. Thus by identifying the module with the highest responsibility signal we can follow the policy developed in that module to solve the problem in an efficient manner. We can extend the model described above to incorporate the modular RL framework. The schematic for the extended model is given in Figure 3.

**Figure 3.**
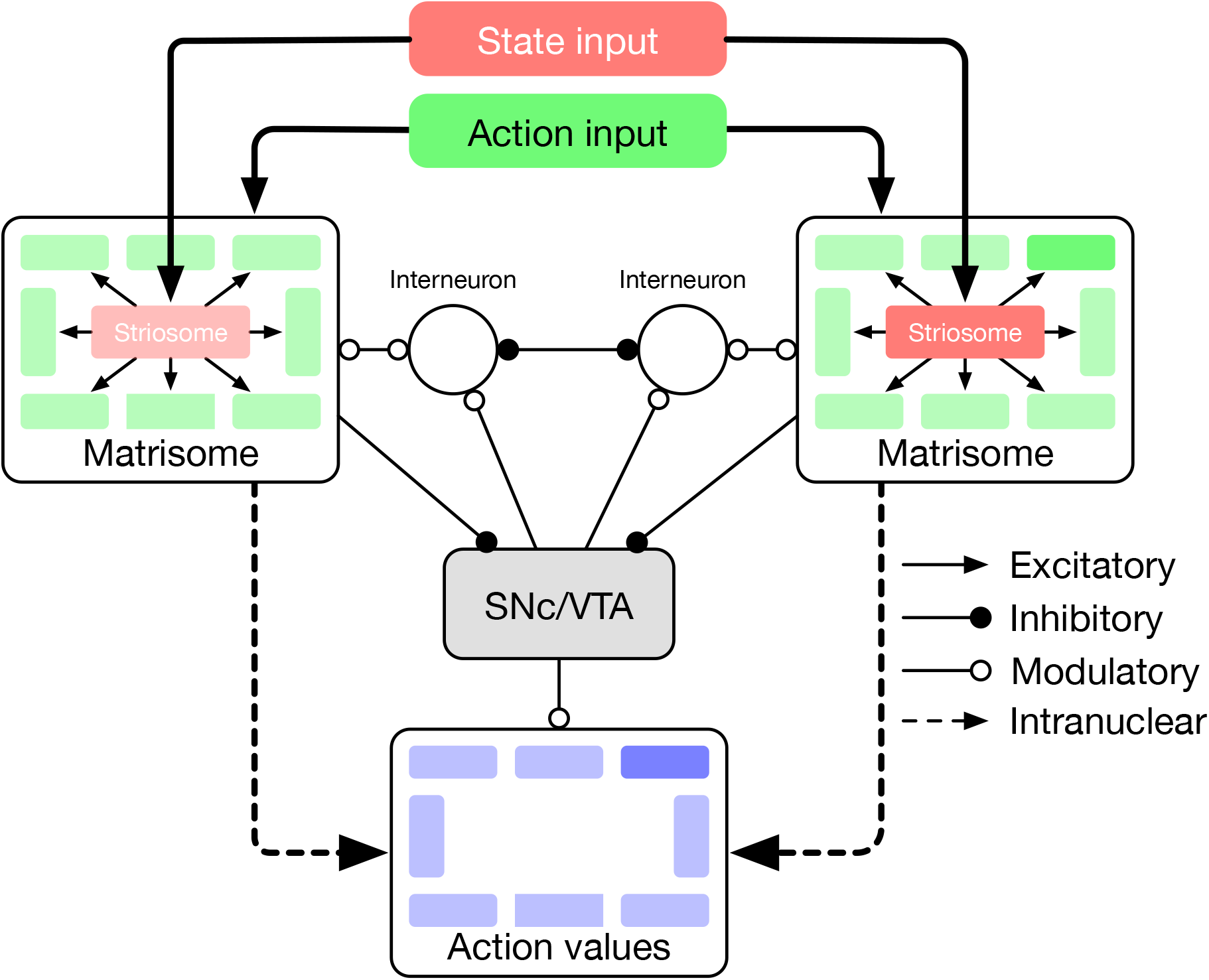
A schematic of the extended model to handle modular RL tasks showing the case with two striatal modules. The state representations of the two modules are used to calculate their respective responsibilities which are then used by the striatal interneurons to choose the appropriate module.

We believe that context selection happens at the level of the striatum and the context modulated activity is projected to the action value neurons. For clarity, we have expanded the intranuclear activity of the striatum in the model schematic (Fig. 3). Supposing there are K modules denoted by *M*_1_, *M*_2_…, *M_K_*. We now define the weights and activities in the previous sections for each module and denote *M_i_* with each term associated with module Mj. Thus, for a module m, the following variables undergo a change in notation:

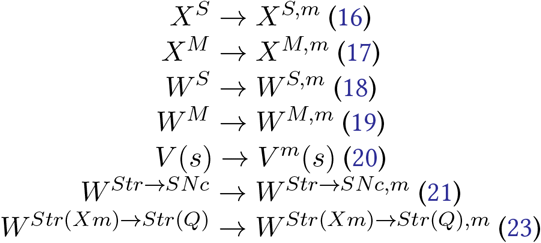

We propose that in addition to the value of the state s, the activity of the Strio-SOM also projects to the SNc to represent the environment feature signal (*ρm*). The weights of these projections are denoted as 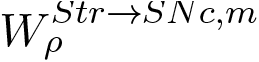 and are trained using the signal from SNc which is representative of context prediction error (*δ\**). The corresponding equations are given in (25) and (26). The context prediction error is calculated as 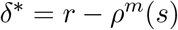

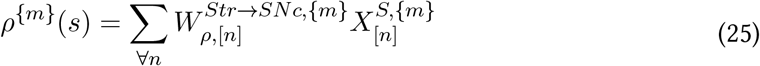

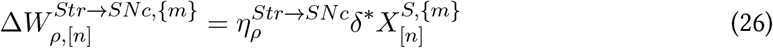

The responsibility signal for each module is denoted by *λ^m^* for module m. In a given state s, the module with the highest λ is chosen for deciding the action in that state. Biologically, we believe that this selection of the appropriate module for the context is guided by the striatal interneurons (Sullivan, Chen, & Morikawa, 2008). Let the winning module in the state s be denoted by *m\**. The winning module projects to the action value neurons (27) following which the processing is the same as in the previous section.

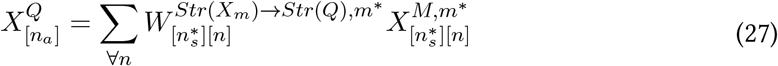

The dynamics of the responsibility signal is given in (28)

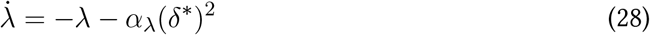

where *α*_λ_ controls the influence of context prediction error on the responsibility signal and *δ\** is the context prediction error.

## 3 Results

### 3.1 T-Maze tasks

The study of non stationary bandit tasks is a reasonably underexplored area owing to the complexity of decision making involved in these tasks. However, some of the earlier results (Lloyd & Leslie, 2013) make some predictions which we aim to replicate with our model. The task performed by the agent is a T-maze task (Olton, 1979) where the agent has to choose one of the arms in a maze. Upon choosing the arm, the agent gets a reward *R_max_* with a given probability (*P_success_*) and a reward *R_min_* with a given probability (*P_failure_*). The task can be extended to a context-dependent problem by reversing the reward distributions with trials.

We study the performance with changing *R_max_/R_min_* and *P_success_/P_failure_*. Animals tend to choose rewards which have a higher magnitude and greater rewards lead to faster convergence (Figure 4A). Similarly, with the same magnitude, animals tend to prefer distributions which reward with a higher probability (Figure 4C). These effects are captured by our model as shown in Figure 4B and Figure 4D respectively. The figures show the ratio of the correct choices by the agent in 50 trials averaged over 50 sessions. The value of exploration factor, p (11) was set as 0.1 and the window length, *τ* (7) was chosen as 5.

**Figure 4.**
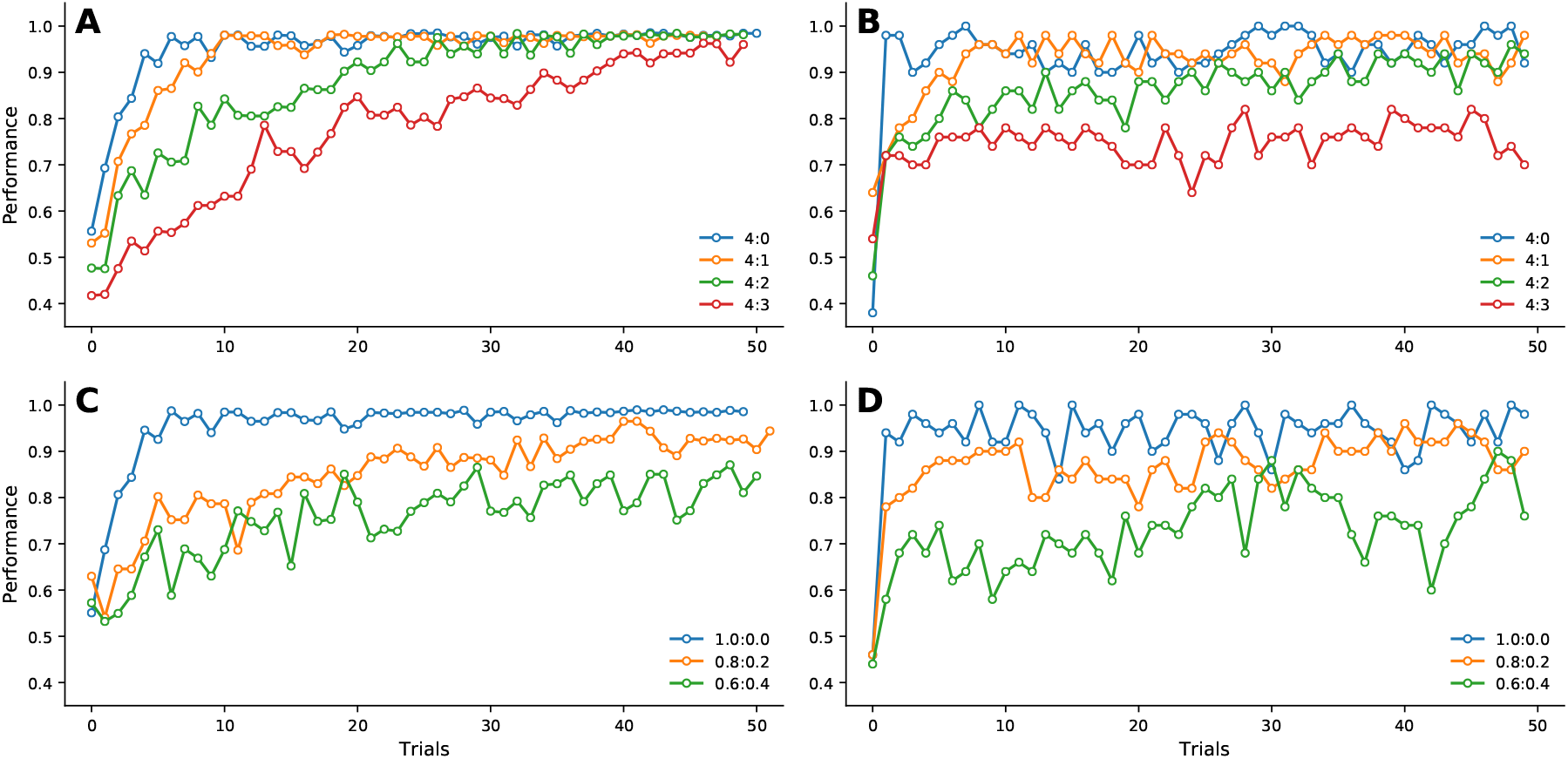
**A**. Results from the (Lloyd & Leslie, 2013) model according to the differences in reward magnitude between the two arms, reward probability being the same. **B**. Results from the proposed neural model. Only the 4:3 ratio case prevents the model from converging. **C**. Results from the (Lloyd & Leslie, 2013) model according to the differences in reward probabilities between the two arms, reward magnitude being the same. **D**. Results from the proposed neural model that are comparable (but slightly worse) to the peformances of (Lloyd & Leslie, 2013).

Experimental evidence (Brunswik, 1939) shows that partial reinforcement and stochastic rewards have a significant effect on reversal learning. We consider a task where the animal is trained on a T-maze with different reward probabilities for 24 trials and then the rewarding probabilities are reversed. We look at the percentage of the trials where the animal chooses the arm which is unprofitable at first and becomes profitable after the reversal. We can observe that the model results (Figure 5) show similar trends to earlier results (Figure 5A). The tasks with deterministic rewards showed quicker reversal as compared to probabilistic rewards that showed slower policy modulation by the agent.

Stochastic reward distributions also have an effect on extinction (Miltenberger, 2011) of a learned policy. To test this, we consider a task where the animal on a T-maze for 24 trials as above. However, the rewards for both arms are set as 0 following the 24 trials and the rate of unlearning is studied. We observe that definite rewarding tasks show faster extinction as compared to the tasks with stochastic rewards (Figure 5C) which is captured by the model (Figure. 5D).

**Figure 5.**
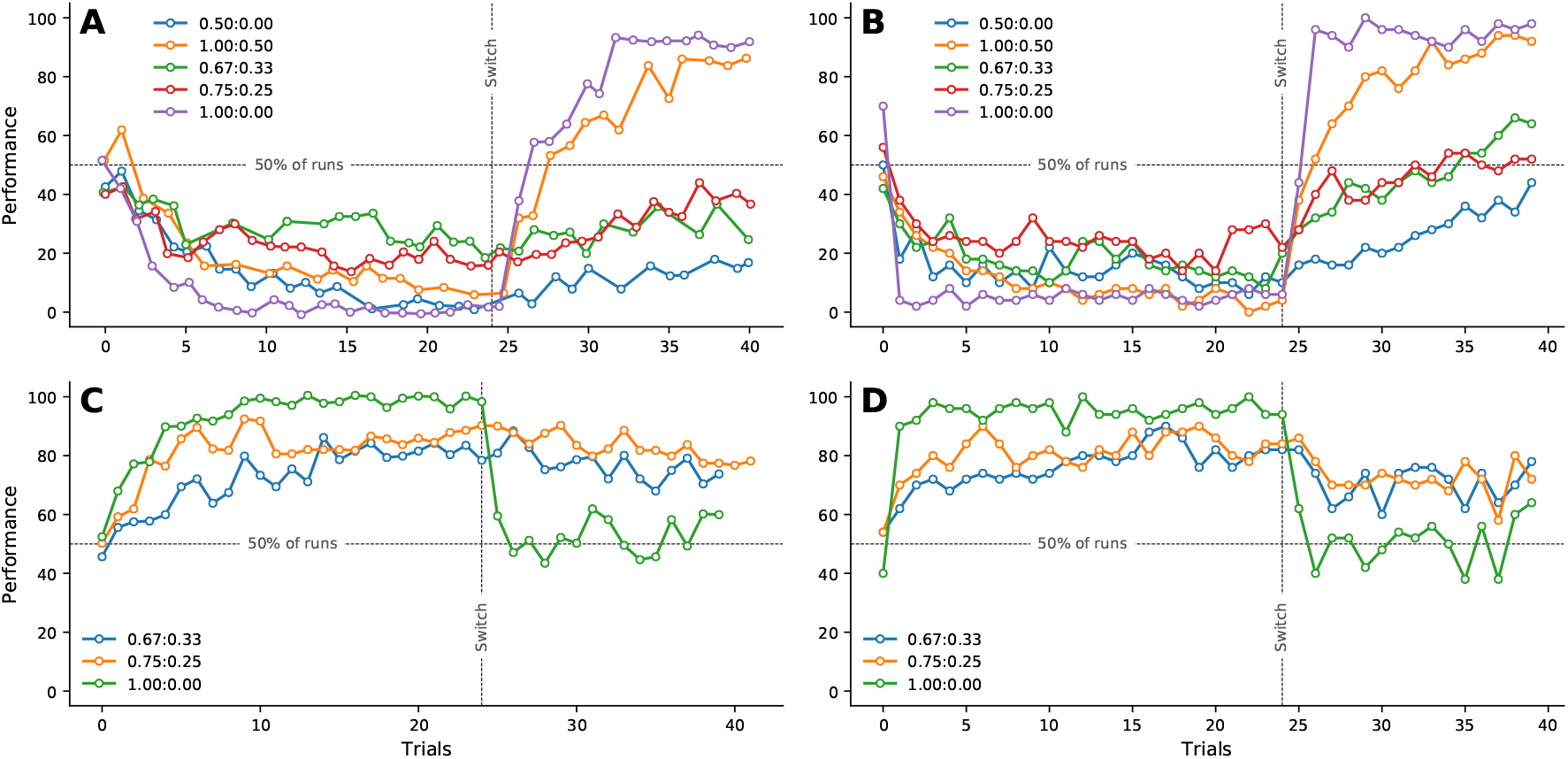
**A**. Results from the (Lloyd & Leslie, 2013) model as the percentage of trials where the animal chooses the arm which is non-profitable for the first 24 trials and becomes profitable following that. **B**. Results from the proposed neural model on the task described in A One can see that the model shows similar trends where the definite reward tasks show faster reversal learning. **C**. Results from the (Lloyd & Leslie, 2013) model as the percentage of trials where the animal chooses the arm which was rewarding before 24 trials following which both arms are not rewarded. **D**. Results from the proposed neural model on the task described in C. One can see that the model shows similar trends as the definite reward task show faster unlearning.

### 3.2 Bandit tasks solved by neural model

In this section, we demonstrate that the proposed model of striatum model is capable of solving bandit tasks. We consider a cue based decision making task where the animal has to choose one of the cues displayed on the screen. This task was first described in (Pasquereau et al., 2007) and a schematic of the task is given in Fig. 6A. The animal is presented with two cues in each trial at two locations (Fig. 6A). Each shape is associated with a different probability of reward. The agent has to choose one of the shapes and gets a reward according to the associated probability. We show that our striatal model is able to solve this task. We consider a 4 dimensional state vector, where each dimension is 1 if the shape is shown and 0 otherwise. The action vector is also 4 dimensional with each dimension denoting the action that is chosen by the agent. The various parameters of the model are given in Table 1.

**Table 1.**
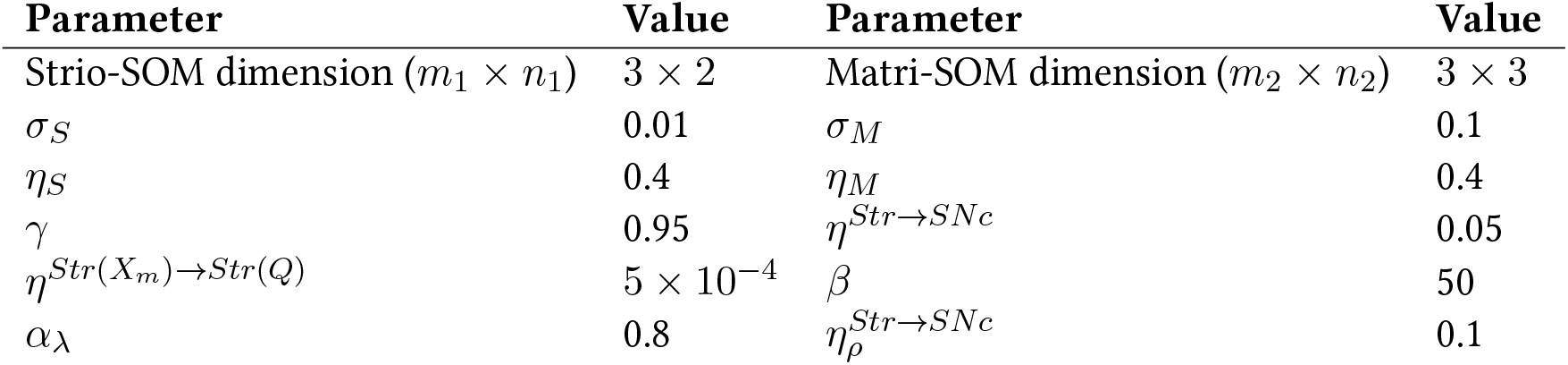
Parameter values for cue based decision making task

The agent (model) is pre-trained where it is given various state and action inputs. We show that the representational maps developed have a center-surround structure (Figure 6C when we view the activity corresponding to all the actions for a particular state. The ratio of correct choices chosen in 200 trials averaged over 25 sessions in given in Figure 6B. Thus, we can see that the agent is able to solve stochastic reward based tasks. Experimental evidence shows that that the percentage of times the agent chooses the arm with reward probability P1, when the ratio of the reward probabilities is P1/(P1+P2), follows a sigmoid activity with center at 0.5 which is well captured by the model (Figure 6D).

**Figure 6.**
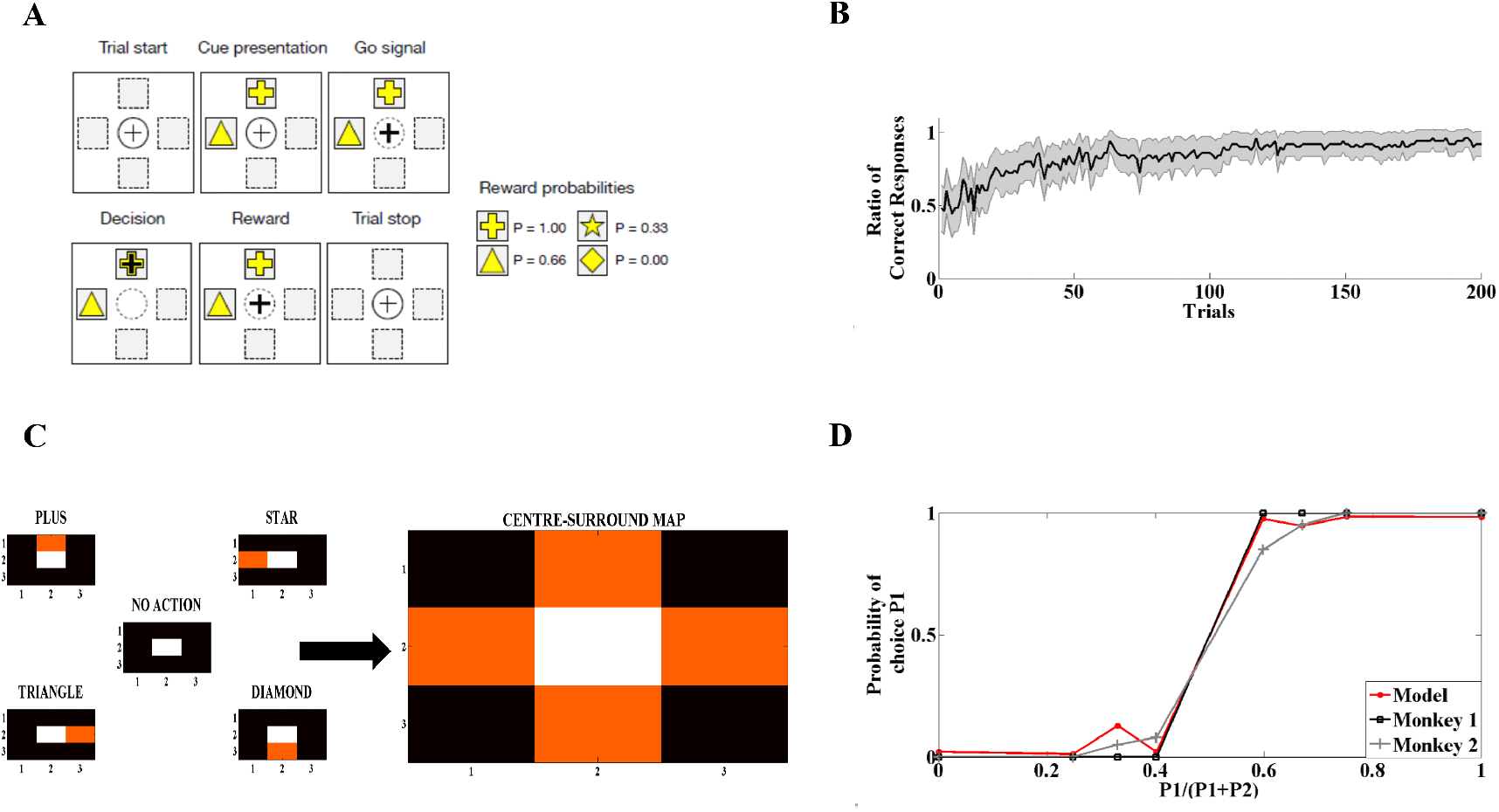
**A**. Schematic of the cue based decision making task where the agent has to choose between the two shapes shown in the screen and each shape has a different probability of reward associated with it. **B**. Percentage of correct responses averaged over 25 sessions for 200 trials. **C**. Mapping of the action inputs forms a center-surround structure when we view the combined activity of the Matri-SOM for all action inputs **D**. Ratio of choosing response 1 with associated probability P1 w.r.t to the sum P1+P2. The model follows a similar trend to the experimental plot adapted from (Pasquereau et al., 2007)

### 3.3 Theoretical vs neural model

We have introduced both a theoretical model capable of solving non stationary bandit tasks and a neural network model which provides a biologically plausible mechanism for the same task. Since there are no available experiments dealing with these tasks (to the best of our knowledge), we shall use the theoretical model to understand the performances of the neural model. In that regard, we use a two arm bandit task which was the underlying problem in both the tasks described beforehand. The reward distributions is reversed after 500 trials and the performance of the agent is characterized by averaging performances over 25 sessions. We also observe the performances for different values of *ϵ* which represents the probability of reward for the non-profitable arm.

Figure 7A (blue) demonstrates the probability of context 1 estimated by the theoretical model whereas Figure 7B (blue) gives the estimation by the neural network model. We observe that the theoretical model is able to identify the context even for larger values of *ϵ*. However, the neural network model is mostly able to identify the context for small values of *ϵ* but fails for larger values. A similar trend can be seen in Figure 7A (orange) and Figure 7B (orange) where we measure the percentage of correct choices by the agent. We observe that the theoretical model is able to learn faster upon context reversal for all values of *ϵ* but the neural model needs to relearn for higher values of *ϵ*.

**Figure 7.**
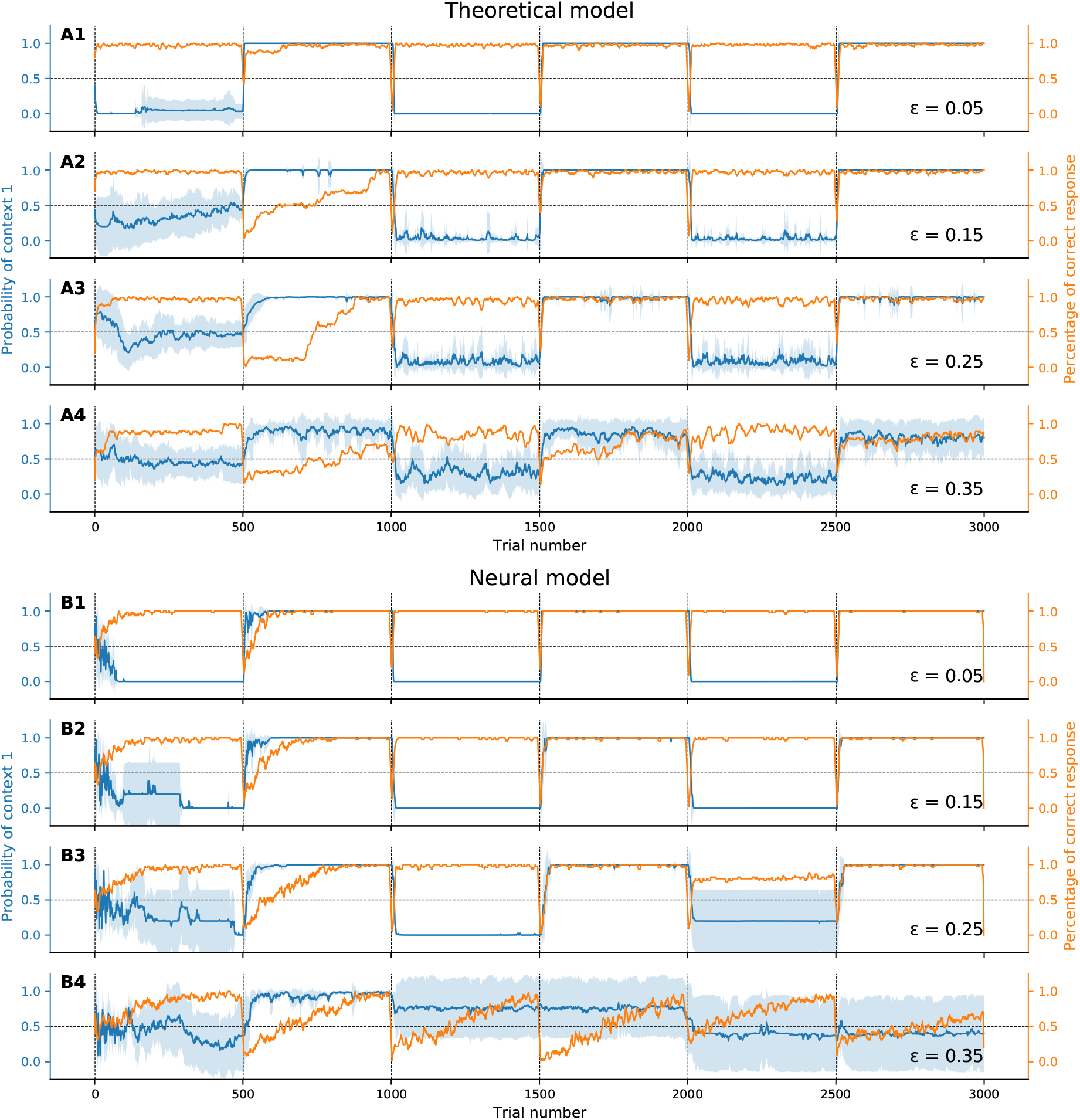
Estimated probability of being in context 1 (blue) and percentage of optimal choices (orange) by the theoretical model, depending on the reward probability *ϵ*. B1 to B4. Estimated probability of being in context 1 (blue) and percentage of optimal choices (orange) by the theoretical model, depending on the reward probability *ϵ*.

From these experimental results, we can conclude that the neural model is able to follow the theoretical model only for low values of *ϵ* and behaves like a single context agent for larger values. This can be further seen in Figure 8 which shows that the neural model performances lies between the theoretical optimal and a single context model and could be the biological mechanism used for solving non stationary bandit tasks.

**Figure 8.**
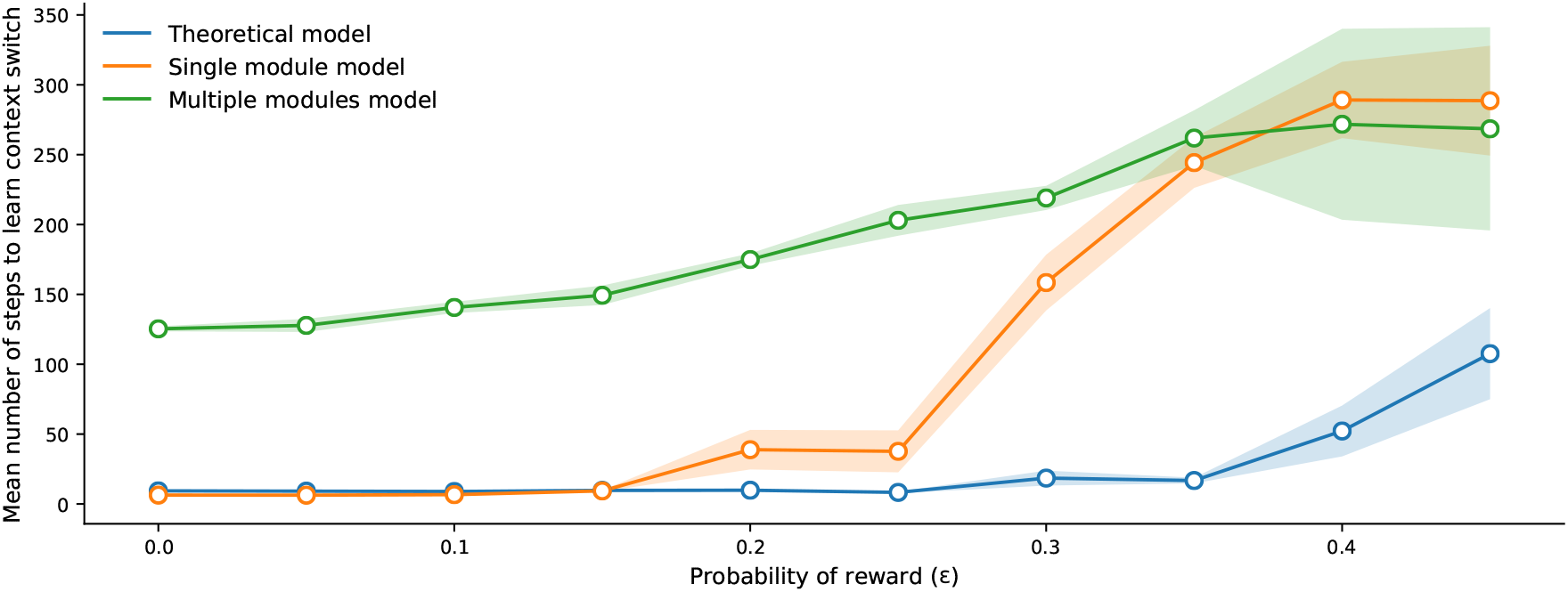
Schematic of the extended model to handle modular RL tasks showing the case with two striatal modules. The state representations of the two modules are used to calculate their respective responsibilities which are then used by the striatal interneurons to choose the appropriate module.

## 4 Discussion

We have presented a theoretical model to solve non stationary bandit tasks. This is also accompanied by a biologically plausible computational model of the striatum which also attempts to tackle these problems.

### 4.1 Adapting to changing contexts

The problem of identifying a change in context in the environment based on the rewards obtained in the previous trials has been extensively studied in the field of change detection (Basseville & Nikiforov, 1993). Given the past history of reward samples upon taking a particular action, Page Hinkley (PH) statistics (Hinkley, 1970) is a popular method for testing the hypothesis that a change in context has occurred (Hartland, Gelly, Baskiotis, Teytaud, & Sebag, 2006; Hartland, Baskiotis, Gelly, Sebag, & Teytaud, 2007). Under the constraint that the rewards come from the exponential family of distributions, PH statistics guarantee minimal expected time before change detection (Lorden, 1971). Our model uses similar ideas of accumulation of mean of rewards in the past trial to predict change in contexts but uses limited memory as a realistic biological constraint. In addition, the model predicts the probability of context change in each trial as opposed to only predicting the occurrence of context change. The model uses information about all the actions in the limited history as opposed to traditional change detection algorithms which assume that the rewards in the history were generated from a single action.

In non stationary bandit tasks, the agent needs to choose an action in each trial in addition to identifying contexts. Earlier studies have shown that sliding window UCB (SW-UCB) (Garivier & Moulines, 2011) is a theoretically optimal policy for such tasks. In addition to being incompatible with biological constraints like limited memory, SW-UCB assumes that the context change is periodic with a period T (used to determine a padding function). The policy in our theoretical model is the same as SW-UCB without any padding functions, since the knowledge of the context period is unavailable to the agent.

### 4.2 Theoretical Model as a Constrained Version of the Full Bayesian Model

The inherent complexity of non stationary bandit problems motivated a Bayesian approach to solve these problems. While the full Bayesian model attempted to give the best possible bound for these tasks, the theoretical model aimed to give a characterization of the expected performance under some realistic constraints such as the ones encountered by an animal solving these tasks. One of the key constraints is the assumption of a limited history. Since the animal has finite memory it can use information from only a small and recent history to guide its decision making (Todd, Niv, & Cohen, 2009). Exploration in action selection is a facet of RL and is also observed in earlier studies (Doya, Samejima, Katagiri, & Kawato, 2002). This is also captured as a constraint in our theoretical model (11).

### 4.3 Striatal Microanatomy and Contextual Learning

Our striatal model is derived from a computational model of the basal ganglia proposed for handling non stationary tasks (Shivkumar et al., 2017). The model is based on the assumption that the striosomes map the state space and the matrisomes map the action space. This is supported from earlier results that the striosomes receive input from the orbitofrontal cortex (Eblen & Graybiel, 1995) known for coding reward related states (Wilson et al., 2014). Anatomical studies also show that striosome medium spiny neurons (MSNs) project directly to SNc (Lanciego, Luquin, & Obeso, 2012) which could compute state values as in our model. Evidence suggests that similar to how projections from the striosomes code for state value, projections from the matrisomes code for action value (Doya et al., 2002). Experimental results show the existence of such neurons in the striatum which code specifically for action value (Samejima, Ueda, Doya, & Kimura, 2005). This is well captured in our model as the Matri-SOM projects to action value neurons in out striatal model. Action selection is done using the softmax policy (24) following the action value computation in the striatum. This policy uses a parameter β which controls the exploration of the agent. We believe that this could be the role of STN, GPe and GPi before action selection is done at the level of the thalamus. This is supported by earlier results which suggest that the underlying stochasticity in the soft-max rule could be achieved indirectly by the chaotic dynamics of the STN-GPe loop (Kalva, Rengaswamy, Chakravarthy, & Gupte, 2012).

### 4.4 Comparing the Theoretical and the Neural Model

The two models proposed in our work were developed and validated independently from each other. However, they share some common features and we can observe that the performance of the neural model falls between the performance of the theoretical model and the neural model with a single module (Figure 8). The theoretical model acts as a lower bound to the performance of the neural model for the given stochasticity in the problem. The neural model is able to achieve performance comparable to the theoretical model for low values of *ϵ* but fails to do so for larger *ϵ* where it becomes similar to a single module system. Thus, we predict that our neural model can explain behavior in stochastic multi context tasks for *ϵ* < 0.3. This also allows us to bound performance of the animal performing such tasks in highly stochastic conditions which is challenging from an experimental perspective due to the large number of trials required. Another feature of our theoretical model is that it is a very simple model with no assumptions on the reward or the context distributions. However, despite its simplistic formulation, the model is quite powerful and can capture all the previous results reasonably well. The modular arrangement of identifying context and using it for task selection is very similar to the proposed striatal model. Thus, the striatal model could be a biologically plausible neural implementation of the theoretical model.

## Acknowledgements

We would like to thank Vignesh Muralidharan for the help in the development of the neural model.

